# Statistical Evidence for Common Ancestry: Testing for Signal in Silent Sites

**DOI:** 10.1101/035915

**Authors:** Martin Bontrager, Bret Larget, Cécile Ané, David Baum

## Abstract

1. The common ancestry of life is supported by an enormous body of evidence and is universally accepted within the scientific community. However, some potential sources of data that can be used to test the thesis of common ancestry have not yet been formally analyzed.
2. We developed a new test of common ancestry based on nucleotide sequences at amino acid invariant sites in aligned homologous protein coding genes. We reasoned that since nucleotide variation at amino acid invariant sites is selectively neutral and, thus, unlikely to be due to convergent evolution, the observation that an amino acid is consistently encoded by the same codon sequence in different species could provide strong evidence of their common ancestry. Our method uses the observed variation in codon sequences at amino acid invariant sites as a test statistic, and compares such variation to that which is expected under three different models of codon frequency under the alternative hypothesis of separate ancestry. We also examine hierarchical structure in the nucleotide sequences at amino acid invariant sites and quantified agreement between trees generated from amino acid sequence and those inferred from the nucleotide sequences at amino acid invariant sites.
3. When these tests are applied to the primate families as a test case, we find that observed nucleotide variation at amino acid invariant sites is considerably lower than nucleotide variation predicted by any model of codon frequency under separate ancestry. Phylogenetic trees generated from amino-acid invariant site nucleotide data agree with those generated from protein-coding data, and there is far more hierarchical structure in amino-acid invariant site data than would be expected under separate ancestry.
4. We definitively reject the separate ancestry of the primate families, and demonstrate that our tests can be applied to any group of interest to test common ancestry.

## Introduction

Charles Darwin laid out the case for the principle of descent from common ancestry (CA) in his *Origin of Species* (1860). Today, descent from CA is widely accepted as fact, a situation which means the underlying evidence is rarely given the full consideration it deserves. While understandable given the strength of the evidence, it is also unfortunate that there have been so few efforts to formally test CA. This scientific vacuum has been filled with misconceptions and half-truths. To amend the problem we must explore the diversity of kinds of data and methods of analysis that can be used to rigorously test CA against the alternative hypothesis of separate ancestry (SA).

The first formal attempt to test CA was undertaken by Penny et al (1982) using phylogenetic trees generated from five different proteins across 11 species of mammals. Their SA model supposed independent origins of each species, with each gene’s optimal tree merely reflecting the outcome of a phylogenetic analysis conducted on data that did not derive from a tree. Under this SA model we would expect proteins to yield rather different phylogenetic trees. Penny et al. (1982) generated a null distribution under SA and used this to show that proteins yielded much more similar trees than would be expected under the SA model.

An underlying assumption of the Penny et al. SA model, which also affects many other statistical tests of common ancestry (Baum et al. 2015), is that there are no functional constraints that might result in proteins derived from SA nonetheless showing a common hierarchical structure. This need not be the case. For example, to rescue the SA hypothesis one might posit that gorillas and chimpanzees share a unique set of functional constraints that explain why all their proteins are more similar to one another than to, say, a lemur. The argument might be that organisms sharing similar form, ecology, behavior and development might share protein sequences that are consistently similar since those proteins are called upon to perform similar functions. Thus, we can conceive a SA hypothesis under which two species that look and behave similarly may be expected to have similar protein sequences due not to common ancestry but to shared protein functions. What is needed to test CA against this particular formulation of the hypothesis of SA is a set of hereditary data that is not susceptible to functional constraints that could result in such artifactual hierarchical structure.

The degeneracy of the genetic code means that a subset of coding sequence variation is silent and thus immune to functional constraint based on protein function. We have known since the 1960s that amino acids are encoded by degenerate, non-overlapping, three-nucleotide codons (Crick et al. 1961). For example, the amino acid valine can be encoded by four different codons (GTA, GTT, GTC, and GTG) differing in their third nucleotide position only *(Figure 1, codon* 3). The precise codon used plays no role in the function of the encoded protein (but, see discussion). Thus, any nucleotide variation at amino acid invariant sites (i.e., sites where the same amino acid occurs in all species) should be free of the kind of functional constraint that could most easily confound statistical tests of SA vs. CA. Thus, finding that different species share the same nucleotide sequences at amino acid invariant positions more often than would be expected by chance can provide strong evidence in favor of CA. These facts allow us to develop statistical tests based on the expectation under SA that codons at homologous amino acid positions should be random and lacking in hierarchical structure.

**Figure 1.**
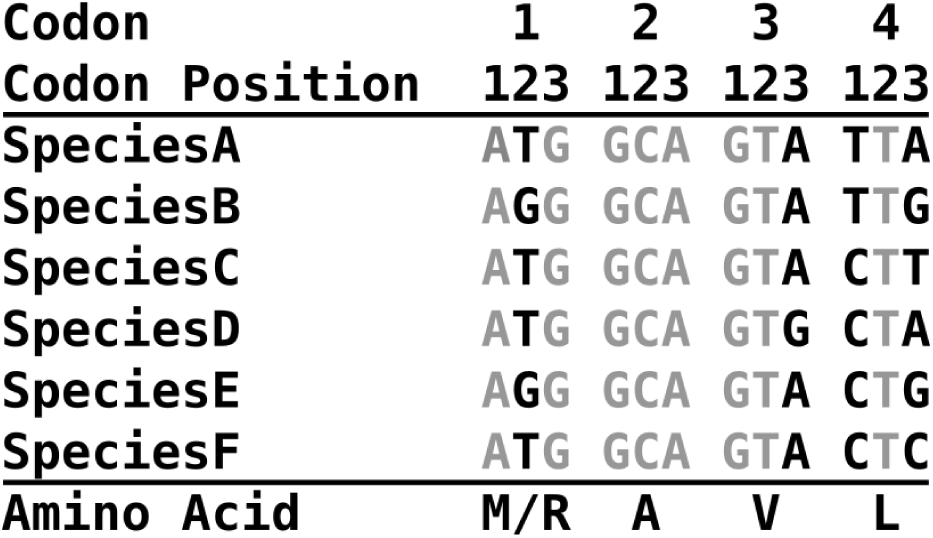
Examples of nucleotide and protein variation. Example DNA sequence from homologous protein-coding genes from 6 different species are aligned in the proper reading frame. Nucleotide positions that vary between taxa are highlighted in black. Codon (1) illustrates nucleotide variation that results in amino acids variation whereas codons (2), (3), and (4) are amino acid invariant. There is no nucleotide variation in codon (2), even though the third position is free to vary and still encode alanine. This is an invariant codon site and is not expected under SA except by chance. Entropy at codon (2) is zero. Codon (3) is variable at the 3rd position, but still codes for valine in all species (entropy = 0.451). There is less variation in codon usage at this site than might be expected under SA since the third position is also free to vary to "T" or "C" and still encode valine. Codon (4) is a six-fold degenerate codon and variable at the first and third position (entropy = 1.79). All potential leucine codons are used.

We set out to test CA vs. SA using information from nucleotide sequences at amino acid invariant sites and a model of SA that posits the independent origin of different taxonomic families (Baum et al. 2015). This family SA model resembles the model laid out by some modern creationists who have proposed that different “types” of lineages, usually equated with families, arose independently but then diversified following the Noahic flood via phylogenetic branching and adaptive evolution to generate the modern diversity of living species (Gishlick 2006). We used the primates as a test case for our examination of CA, in part because the common ancestry of all primates, including humans, remains controversial among some skeptics of CA. Additionally, the primates are a young enough clade that we might reasonably expect some phylogenetic signal even in rapidly evolving sites such as degenerate third codon positions.

Generating expectations under SA for the frequency with which two taxa share the same codon at a given amino acid invariant site is not as straightforward as simply assigning each possible codon an equalchance of use. There are well known patterns of codon usage bias genome-wide related to variability in GC content across the genome (Plotkin and Kudla 2011), which need to be taken into account in our null model. We developed three different expected codon usage distributions at invariant amino acid positions under SA (see Materials and Methods). We also modified the tests to mask positions in which the exact same codon was used in all taxa, since such invariant codon usage at a position might be due to some unknown functional constraint *(Figure 1, codon* 2).

In addition to testing the hypothesis that codons observed at invariant amino acid sites are less variable across families than would be expected under SA, we considered two other patterns that have been proposed to provide evidence of CA. First, following Penny et al. (1982) we assessed whether the topology of trees inferred from *silent sites*, that is from nucleotide variation at amino acid invariant sites, agree with trees inferred from the variable amino acid sites more than expected by chance. Finally, we conducted permutation tail probability (PTP) tests (Archie 1989; Faith 1989) on nucleotide variation at amino acid invariant sites to see if their internal hierarchical structure is greater than expected under SA. All three statistical approaches roundly rejected family SA in favor of CA. In combination with recent companion papers which use other novel methods to test primate CA (Baum et al. 2015; Larget et al. 2015), these results statistically negate the hypothesis of SA of the primate families.

## Results

### Shared codon usage

We studied 17 coding genes comprising a total of 3657 aligned codons. We generated 50 datasets, each of which included one representative species per family for each gene. Across these 50 data sets, the number of amino-acid invariant sites ranged from 1742 to 2172 (ignoring sites encoding methionine and tryptophan which lack codon degeneracy). Of the amino acid invariant sites, 823 to 1045 (~48%) had some variation at the nucleotide level.

For each of the 50 data sets, we calculated the observed summed entropy value for the amino acid invariant sites and also the expected value under each of the three codon-usage models. We also repeated these calculations after dropping those codons without any silent variation. As summarized in *Figure 2*, across all six conditions, the observed data had much lower entropy than expected under the family SA hypothesis. The number of standard deviations between paired observed and expected entropy statistics ranged from 85.8 to 313.1 across all six conditions. These *z*-scores correspond to p-values effectively equal to zero and represent enormously strong rejection of the family SA model.

**Figure 2.**
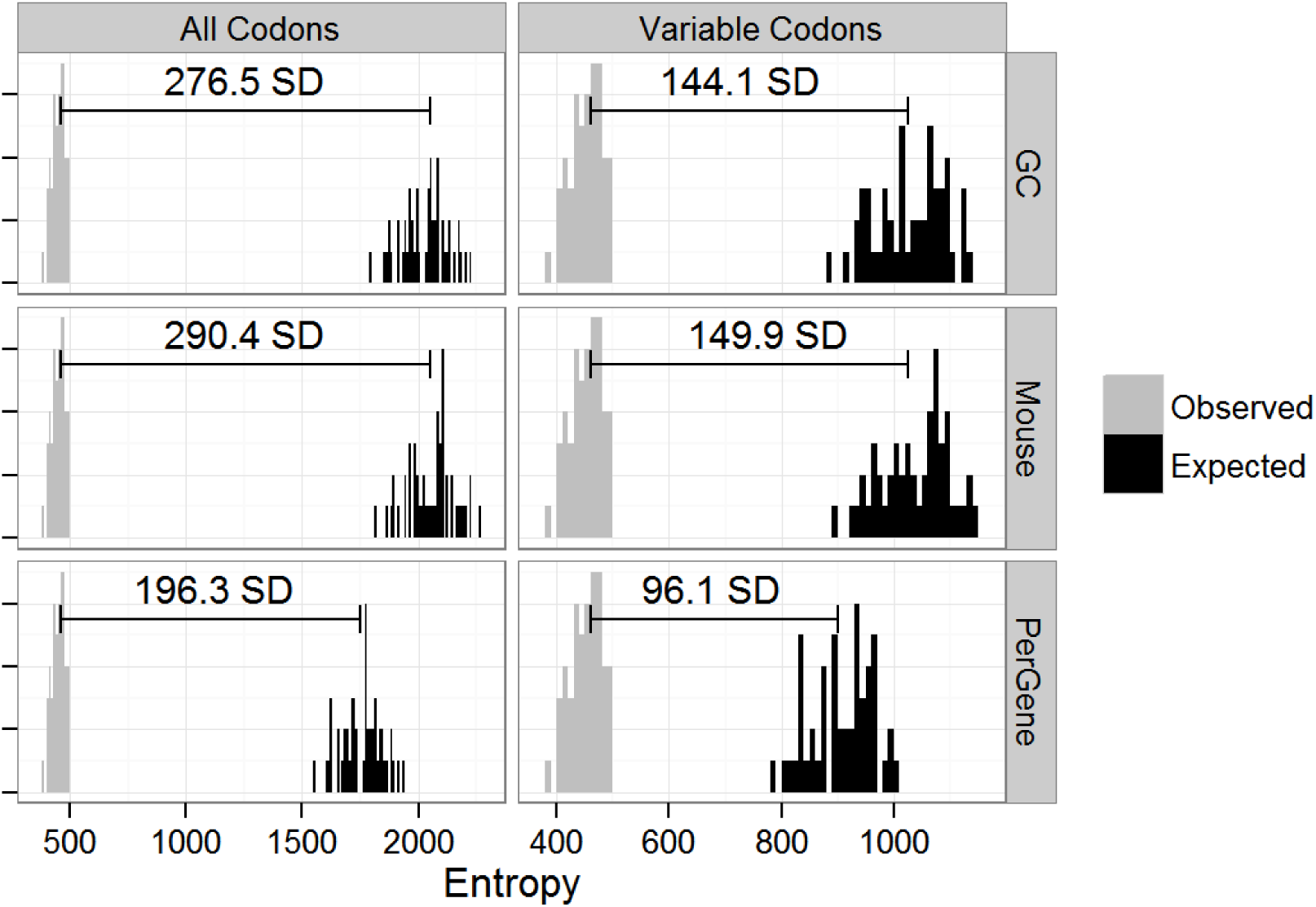
Observed and Expected Entropy in Codon Usage at Amino Acid Invariant Positions under Different Models –. Histograms show the observed and mean expected entropy for 50 data sets, each composed of a different sampling of species to represent each gene for each family. The horizontal bars represent the mean distance (across the 50 data sets) between the observed and expected values in units of standard distribution (SD). Expected entropy values were derived from alignments generated from one of three models of codon usage (“GC”, “Mouse”, and “PerGene”) and for each model either included all amino acid invariant sites (“All Codons”) or only amino acid invariant sites with some variation in nucleotide sequence (“Variable Codons.”)

As we would expect, rejection of SA was strongest when we included all amino acid invariant sites rather than just those that show some nucleotide variation among families. This is because the entropy statistic at nucleotide invariant codon sites is 0, and all codon distribution models yield an expected entropy greater than zero. Using a codon null model that considers GC content yielded expected entropy values that were somewhat closer to the observed value than using the mouse whole-genome codon usage. Using a model based on per-gene codon usage resulted in even lower distances between observed and expected values, confirming our expectation that this would be the most conservative test.

Most of the variation in observed and expected entropy values across the 50 data sets was due to variation in the length of the sequences of the species chosen to represent a family. Observed and expected entropy both increase in longer alignments, as expected, but we also observed that as the length of the alignment increased, so did the z-score *(Figure 3)*. This suggests that adding more sequence data at more coding genes would only serve to strengthen the conclusion that there is significantly less nucleotide variation at amino acid invariant sites than would be expected under primate family SA.

**Figure 3.**
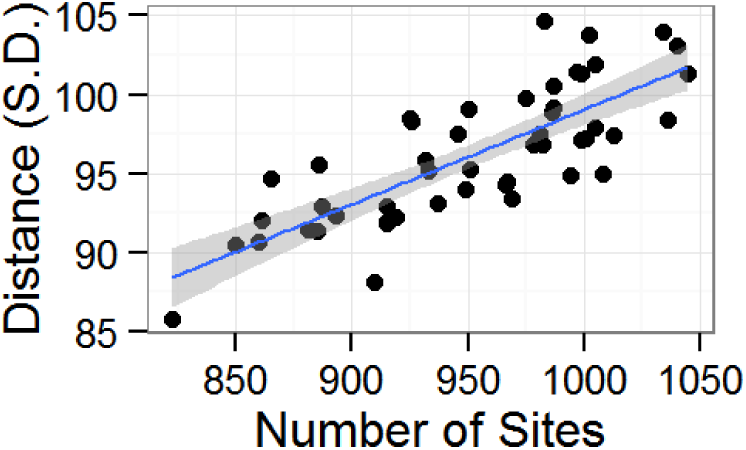
Statistical Strength vs. Alignment Length –. As the number of conserved amino acid sites in the alignment increases, so does the statistical strength of rejection of SA. If we were to include more primate gene sequences in these alignments this suggests that the strength support for CA would only grow. The grey bar indicates 95% C.I. for the regression line.

## Tree agreement

The Robinson-Foulds distance between trees estimated from nucleotide variation at amino acid invariant sites and from amino acid variation ranged from 2–10 across the 50 data sets (mean =5.6, 39 replicates had RF distance = 4 or 6). Using the exact p-values for significance from table 4 in Hendy et al. (1984) for 16-taxon trees, the largest p-value across all 50 replicates was 3.12 × 10^−9^, the median p-value was 2.48 × 10^-11^, and the smallest was 1.27 × 10^-13^. These results show that the two trees generated from nucleotide sequence at amino acid invariant sites and variable amino acid sequence are significantly more similar to one another than trees drawn at random from the full tree space. This is readily explained if the gene sequences are related by descent along the primate phylogeny. However, this result would not be expected under SA, which would suggest that the trees estimated are due to shared functional constrains, since there is no reason to expect functional constraints to be the same at the amino acid and nucleotide levels.

### Permutation test

The permutation tail probability test on nucleotide variation at amino acid invariant sites showed that the original unpermuted data has much more hierarchical structure than would be expected by chance under SA (Figure 4). The optimal trees for the original data differed in length from the permuted data by an average of more than 100 standard deviations (of the permuted data), corresponding to a p-value effectively equal to zero. This result also provides compelling support for the CA model of family SA.

**Figure 4.**
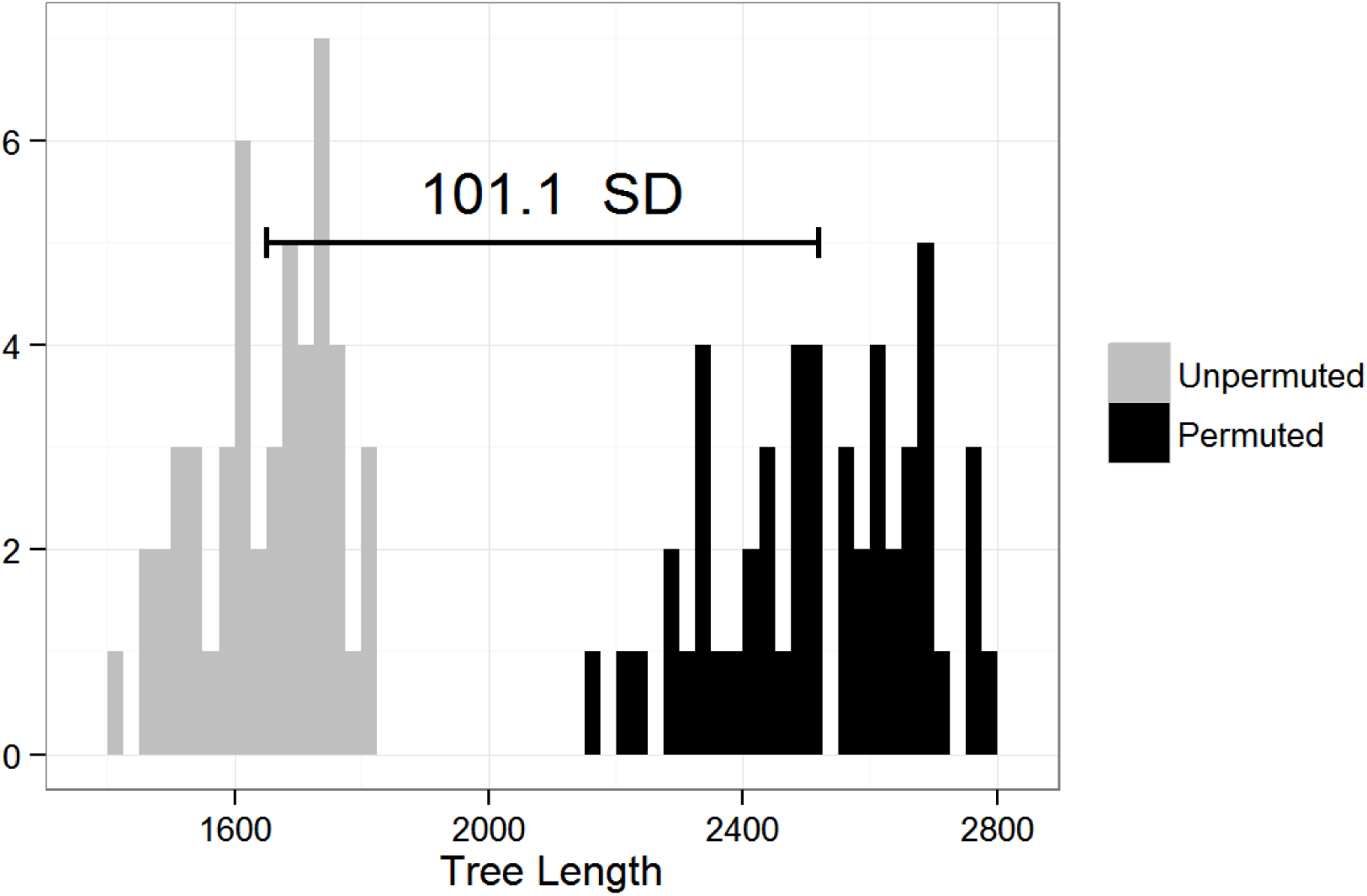
PTP test –. For each of the 50 trials, the conserved amino acid sites underlying DNA sequence were used to build a phylogenetic tree. The tree length of the original unpermuted data (light grey) is paired with a tree length distribution on permuted data. Tree lengths of unpermuted data are on average 101.1 standard deviations from their paired permuted tree lengths

## Discussion

Testing the hypothesis of SA using codon variation at amino-acid invariant sites is heavily dependent on the model of codon usage expected under SA. We have suggested three such models, and tested the hypothesis of SA under each. The mouse whole-genome codon usage model effectively captures gross patterns of codon usage shared across mammalian genomes. The per-gene codon-usage distribution might be the most accurate if we had access to the full coding sequence of all genes in this study. However, the short length and complete absence of certain codons in some genes artificially skews the expected entropy value lower (i.e. towards being more favorable to SA). The GC-content model is probably the most biologically realistic model, although our rather simplistic formulation could certainly be improved upon by updating codon usage based on dinucleotide frequencies (Nussinov 1981; Beutler et al. 1989; Antezana and Kreitman 1999). In any case, under all models that we considered, the observed variation across taxa in codon usage at amino acid invariant sites is far less than we would expect under SA, thus strongly supporting CA of the primate families.

Our formulation of the SA model assumes that nucleotide variation in primates at amino-acid invariant sites has little to no functional significance. The question arises as to whether it is the case that such silent variation is truly non-functional and whether alternative codons at amino acid invariant sites produce equivalent protein products with the identical efficiency. It is certainly the case that codons are used unequally in coding genes across the tree of life, a phenomenon called codon bias (reviewed in Plotkin and Kudla 2011). Two classes of explanations have been put forward to explain codon bias: (1) a selective model in which selection operates on translational efficiency and accuracy based on the relative abundances of different tRNA classes and (2) a neutral model in which mutational processes result in codon bias (see Sharp et al. 1995; Hershberg and Petrov 2008 for reviews). In some taxa there is evidence for selective pressure on codon usage via translational efficiency and protein structure. Specifically, the commonest major codons are translated more rapidly, with greater fidelity, and in ways that promote proper folding when compared to the minor codons (Bennetzen and Hall 1982; Ikemura 1985; Sharp and Cowe 1991; Moriyama and Hartl 1993; Akashi 1994). There is some evidence of selective pressure on codon usage in vertebrates (Chamary et al. 2006), but the vast majority of the data suggests that selection on nearly all synonymous positions in vertebrates is weak to non-existent (Yang and Nielsen 2008). As a result, even if codon bias could lower the expected entropy of independently evolved primate sequences, this effect is likely to be minor compared to the great difference between the observed and expected entropy values that we detected under all models (even after excluding invariant codons, which would presumably be most easily explained by convergent evolution due to selection on codon bias).

Supposing for a moment that there is an alternative model of codon usage at degenerate positions under which each separately evolved clade would tend to share similar nucleotides at amino-acid invariant sites, this fact would not explain the results of the tree agreement and PTP tests. Whereas codon bias could affect protein translation rate and folding, the primary amino acid sequence might affect protein function, which is unlikely to result in a similar hierarchical pattern of variation. Thus, even a very high level of codon bias would not lead to the expectation under SA that a tree inferred from silent codon variation would resemble that inferred from amino acid sequences. Likewise, even a strong and specific codon bias would still predict that nucleotide variability at amino-acid invariant positions would be randomly distributed among lineages. But, this is contradicted by PTP tests which show that different amino-acid invariant sites have much more similar pattern of silent nucleotide variation than expected under SA. Thus, our entropy analyses, tree-agreement tests, and PTP tests provide independent and consistently overwhelmingly strong rejection of the hypothesis of separate ancestry for primate families.

These tests on entropy and hierarchical signal at amino-acid invariant sites can be applied to almost any group of interest. However, through the passage of time, the practicality of these tests will be compromised due to the loss of information (Sober and Steel 2002). Fossil evidence and molecular clock data suggest that the two major suborders of extant primates, the Strepsirrhini and Haplorhini, diverged from a common ancestor between 63 and 87 MYA (Chatterjee et al. 2009; Perelman et al. 2011; Springer et al. 2012). Even given this time since divergence, nucleotide variation at amino-acid invariant sites still shows high levels of phylogenetic signal. We would, thus, expect the tests to also work on other clades of a similar age. However, as one goes further back in time, patterns of similarity will inevitably break down and we would expect silent site variation to converge to that expected under SA. Given such a loss of information, the codon usage test is not useful for testing the universal common ancestry (UCA) of all life on earth, which is well known asa difficult problem (Theobald 2010; Koonin and Wolf 2010; Yonezawa and Hasegawa 2010, 2012; de Oliveira Martins and Posada 2014). Nonetheless, as applied to a relatively young group like the primates, the tests described here provide enormous statistical support to the CA hypothesis such that anyone who was previously unconvinced of primate CA, and was open to statistical evidence, should be entirely convinced that primate families did not originate independently.

## Materials and Methods

### Shared nucleotide variation at amino-acid invariant sites

Sequences from 54 primate genes in 84–186 primate species generated by Perelman et al. (2011) were obtained through NCBI GenBank. For each gene, sequences were arranged in a multiple alignment with MUSCLE (Edgar 2004). We aligned the longest isoform human mRNA sequences of the corresponding genes to the full primate multiple alignments. The 37 sequences that included little-to-no coding sequence were discarded, resulting in data sets composed of the coding sequence from 17 genes. We trimmed the multiple alignments so as to reflect the correct reading frame. For each alignment studied, we randomly picked the protein-coding sequence from one species in each of the 16 families for which there was sequence data as the family representative sequence and we repeated this procedure 50 times. This resulted in 50 different data sets each containing 17 genes and one representative species for each the 16 primate families, but with different species potentially representing each family and each gene.

At each codon in the alignments for which the encoded amino acid was identical across all included species (amino acid invariant sites), and at which there was the possibility for variable codon usage due to codon degeneracy, the observed variation in codon usage was measured via the entropy at that position. Entropy quantifies the amount of variability in a data set where values are categorical. Our formulation of the entropy statistic (*H*) at each position in the protein alignment was:

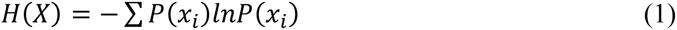

where *x_i_* is an observed codon at position *i* and *P*(*x_i_*) is the frequency of codon *x_i_* observed at that position. For each of the 50 datasets, we summed the entropy across every position in all 17 protein sequences to generate a single observed entropy statistic.

In order for our summed entropy statistic to be useful for testing CA vs. SA, we needed to generate a null distribution under family SA. We developed three different expected codon usage distributions and compared the summed observed entropy statistic to entropy values expected under these null distributions. To do this we used random draws from an expected codon-usage frequency distribution to pick a codon at each invariant amino acid position in the alignment. If, for example, all 16 family representatives encoded valine at a given invariant amino acid position, we used one of three codon usage frequency tables to draw 16 expected valine codons at that position. From these newly generated DNA sequence alignments we calculated the summed entropy value and repeated this procedure 100 times, calculating the expected mean entropy and its standard deviation under each.

Under the first, and most general expected distribution, we assumed that primate codon usage resembles that which is observed across the whole genome of the house mouse *Mus musculus*. Differential usage of codons, codon bias, is remarkably consistent across mammals according to codon usage in annotated reference genomes in the NCBI database. The mouse genome-wide codon usage frequency for valine, for example, is 0.17, 0.12, 0.25, and 0.46 for the codons GTT, GTA, GTC, and GTG respectively, which is very close to that seen in *Homo sapiens:* 0.18, 0.11, 0.24, and 0.47. Although codon usage patterns for each amino acid are available for the full human genome, we considered it more appropriate to use a mammalian relative instead of an extant primate to avoid any potential bias from using one extant primate to generate expectations for all others. In practice, however, genome-wide human and mouse codon usage distributions are extraordinarily similar and both yield essentially the same results (not shown).

A second codon usage distribution was generated on a per-gene basis, based on GC content at degenerate positions. Analysis of codon usage across taxa suggests that GC content of adjacent non-coding DNA and degenerate positions is a better predictor of codon usage in a given gene than is genome-wide codon usage (Matassi et al. 1999; Knight et al. 2001). We calculated the GC content (GC%) for each gene based only on degenerate codon positions in that gene (in any species) in that data set. To predict codon usage frequency under independent SA, we assumed that the frequencies of A and T at degenerate positions are equal and, likewise, that the frequencies of G and C are equal. For example if the GC content in degenerate codon positions of a gene was 60%, for the amino acid valine we would set the expected codon usage to 0.2, 0.2, 0.4, and 0.4 for GTT, GTA, GTC, and GTG respectively. Although the true picture of codon usage is more complicated (Nussinov 1981; Beutler et al. 1989; Zhang et al. 1991), including this approach as one of three different models is a reasonable approximation.

Finally, and perhaps most conservatively (as it is least likely *a priori* to reject SA), we calculated expected codon usage for each amino acid based on that amino acid’s observed codon usage across all species included in the analysis. In this case we used the entire coding alignment (84–186 species) and all positions including variable amino acid positions and calculated the observed frequency of each codon in each gene. For example, if the observed relative frequency of the four valine codons in the complete alignment for a gene was 0.1, 0.2, 0.4, 0.3, for GTT, GTA, GTC, and GTG then this would be selected as the expected frequency for each amino acid invariant site encoding valine in this gene.

For each of these codon usage distributions we generated two summed entropy values: one in which we included all amino acid invariant positions and the other in which we included only amino acid invariant positions that showed *some* nucleotide variability among the sampled species. Thus, an expected entropy distribution was calculated for six conditions: mouse codon usage, GC codon usage, and per-gene codon usage at both all amino acid invariant positions and at a subset of amino acid invariant positions that showed nucleotide variation. We then calculated the distance between the observed entropy statistic and the expected entropy distribution in terms of standard deviations from the expected entropy mean. All scripts and aligned input data are available online at https://github.com/mbontrager/primate_ca.

### Tree agreement

We set out to test whether the agreement among the optimal trees inferred from silent sites with those inferred from amino acid sites is greater than expected by chance. We conducted this test for 50 data sets each composed of one representative species for each of the 16 primate families, with the family representatives chosen randomly from the Perelman et al. (2011) data set (see above). In each data set, we identified amino acid invariant sites and retained their corresponding nucleotides to form a silent-site nucleotide alignment. We assessed whether the tree inferred from 50 silent-site alignments differed significantly from the tree inferred from the 50 corresponding amino acid sequences. Even if there were constraints on protein function that imposed hierarchical structure on amino acids, barring CA there is no reason to expect a similar hierarchical structure in silent sites (see discussion).

The method we used was described in more detail in Baum et al. (2015). Briefly, we estimated gene trees using maximum likelihood with the program RAxML version 8.0.24 (Stamatakis 2006). For the DNA silent-site alignments we estimated trees with the GTR+CAT model and 10 distinct starting trees for optimization. For amino acid alignments we used automatic model selection plus gamma-distributed rates using BIC on the most parsimonious tree and 4 distinct starting trees. For each of the 50 replicates we quantified the tree-to tree similarity between silent-site and coding-site trees using the Robinson-Foulds (RF) distance (Robinson and Foulds 1981). These calculations were done in R (R Core Team 2014). The expected tree-to-tree similarity under SA is predicted from the set of all random trees with 16 taxa, as determined by (Hendy et al. 1984)(table 4, p.1063). This allows us to look at the optimal trees from two data sets and calculate a p-value: the probability of observing as great or greater similarity if the trees were drawn independently at random from the space of all trees.

### Permutation test for hierarchical structure

One signature of CA is the presence of hierarchical structure within a sequence alignment, which results from each position in the alignment having undergone mutation along the same branching gene tree. In the case of SA, in contrast, we should expect different sites in a sequence to vary independently, which would tend to be manifested by the data set requiring abundant homoplasy when mapped onto its optimal phylogenetic tree. The permutation tail probability test (Archie 1989; Faith 1989) assesses hierarchical structure within a data set, and thus provides a means to test whether the data are consistent with SA (Baum & Smith, 2013; Baum et al. 2015). We analyzed the 50 silent site alignments to see if they contain more internal phylogenetic structure than expected by chance.

We conducted the PTP test in a parsimony context, using tree length as our measure of fit between the data set and its optimal tree. The 50 data sets were analyzed in PAUP* 4.0a137 and a PTP test was run using the permute command with 1000 replicates, each using simple addition sequence, TBR heuristic searches with maxtrees set to 100. The mean and standard deviation of the permuted data sets was determined and was used to calculate a p-value for the observed (unpermuted) data set using a normal distribution.

## Acknowledgements

The authors would like to acknowledge the University of Wisconsin-Madison Department of Botany, and the participants in the Botany seminar class (Botany 940) who helped to develop the ideas for this paper. In particular, we would like to thank Peggy Boone, Chloe Drummond, Scott Hartman, Lam Si Tung Ho, Steven Hunter, Michael Johnson, Bill Saucier, and Claudia Solis-Lemus for their contributions during the seminar class. We would also like to thank Elliott Sober and Mike Steel for their feedback on the methods presented above during the seminar class.

